# Behavioural analysis of swarming mosquitoes reveals higher hearing sensitivity than previously measured with electrophysiology methods

**DOI:** 10.1101/2021.09.13.460025

**Authors:** Lionel Feugère, Olivier Roux, Gabriella Gibson

## Abstract

Mosquitoes of many species mate in station-keeping swarms. Mating chases ensue as soon as a male detects the flight tones of a female with his auditory organs. Previous studies of hearing thresholds have mainly used electrophysiological methods that prevent the mosquito from flying naturally. The main aim of this study was to quantify behaviourally the sound-level threshold at which males can hear females. Free-flying male *Anopheles coluzzii* were released in a large arena (~2 m high × 2 m × 1 m) with a conspicuous object on the ground that stimulates swarming behaviour. Males were exposed to a range of natural and synthetic played-back sounds of female flight. We monitored the responses of males and their distance to the speaker by recording changes in their wingbeat frequency and angular speed. We show that the mean male behavioural threshold of particle-velocity hearing lies between 13-20 dB SVL (95%-CI). A conservative estimate of 20 dB SVL (i.e., < 0.5 μm/s particle velocity) is already 12 to 26 dB lower than most of the published electrophysiological measurements from the Johnston’s organ. In addition, we suggest that 1) the first harmonic of female flight-sound is sufficient for males to detect her presence, 2) males respond with a greater amplitude to single-female sounds than to the sound of a group of females and 3) the response of males to the playback of the flight sound of a live female is the same as that of a recorded sound of constant frequency and amplitude.

## MAIN TEXT

### Introduction

Hearing is a key sensory modality for mosquito mating; it enables males to detect females at a distance through the combined sounds of their respective flapping wings (Warren et al., 2009; Simões et al., 2018; Feugère et al., 2021b). The more sensitive males are to flight sounds, the further away they can hear a female and the sooner they detect and close in on a nearby female in the context of highly competitive mating-swarms. The male antennal organs of mosquitoes are the most sensitive to sound described so far among arthropods (Göpfert and Robert, 2000), however, the measurement of hearing sensitivity is usually performed on tethered males, which prevents natural body movement such as antennal orientation and wing flapping behaviour in response to female sound. Only a few studies have measured hearing thresholds behaviourally (Menda et al., 2019; Lapshin and Vorontsov, 2021; Feugère et al., 2021b). The measurement of behavioural sound-sensitivity in flying male mosquitoes faces the difficulty of monitoring how much sound energy actually reaches their antennae because the sound level meter is at a fixed-position, whereas the position of the male mosquito is continuously changing during his flight. The aim of this study was to quantify behaviourally the overall sound-level threshold at which males can hear females, i.e. the limit of sensitivity of a male to locate a female in flight. Accordingly, we had to determine the components of female-wingbeat sound that male mosquitoes are most responsive to, so that our definition of the sound level includes only the frequency bands audible to males.

Mosquitoes hear airborne sound by detecting air-particle velocity through friction between air particles and the mosquito’s fibrillae located on the flagellum of their antennae. Unfortunately, there are no instruments that can truly measure particle-velocity on the market as yet (Zhou and Miles, 2017), however, it can be estimated by using pressure-gradient microphones (commonly called ‘particle-velocity microphones’). Another strategy to estimate particle-velocity is to use pressure microphones located in the far-field of the sound source, i.e., where the sound pressure level (SPL) can be approximated to that of sound particle velocity level (SVL). However, SPL hearing thresholds have sometimes been measured under the near-field condition instead of far-field (Tischner, 1953; Belton, 1961; Dou et al., 2021), which means there is a risk that some reported hearing thresholds may have been under-estimated, as elaborated in the Discussion section.

Hearing thresholds can be assessed by measuring a physiological or behavioural response to a given stimulus sound level and sound frequency. Among the physiological methods, laser vibrometry records the vibration of the flagellum (Göpfert et al., 1999; Pennetier et al., 2010), however, it is limited when assessing hearing threshold because the recorded vibration only refers to the input to the hearing chain (i.e., flagella movement) and does not provide any indication as to whether or not the neurons of the mosquito have been neuro-electrically activated following the sound-induced vibration of the flagella. Unlike laser vibrometry, electrical responses of the JOs to airborne sound stimuli result from the complete sensory chain of the auditory system (i.e., from the mechanical vibration of the flagella to the electrical response of the JOs). With this method, the electrical response-threshold in male *Culex pipiens pipiens* JOs showed a mean sensitivity of 32 dB SVL per JO scolopidia (range of 22-44 dB SVL; n=74 JO scolopidia; criterion = 2 dB above noise floor; 18-21°C) (Lapshin and Vorontsov, 2019) and a mean of 44 dB SVL per mosquito in three male *Culex quinquefasciatus* JOs (range of 36-52 dB SVL; n=3 males; criterion = 10 dB above noise floor) (Warren et al., 2009). In *Aedes aegypti,* the male JO nerve was shown to respond to a mean of 40 dB SVL (range of 31-50 dB SVL; n=11 males) (Menda et al., 2019). In some species, such as *Anopheles coluzzii*, the antennal fibrillae are extended only during their active phase, which improves their JO hearing sensitivity by 17 dB in terms of SVL (Pennetier et al., 2010). Under this antennal physiological state, Pennetier *et al.* (2010) measured a JO response-threshold in two male *An. coluzzii* of only 10 dB SVL (range of 5-12 dB SVL, i.e., particle velocity of 1.5±0.6 10^−7^ m/s; n=4 measurements on 2 males; criterion = 1.4 recording noise floor).

In a distortion-product based hearing system, as proposed for mosquitoes, hearing sensitivity can be further enhanced (or even produced) by the mosquito beating its wings (Lapshin, 2012). However, electrophysiological and laser vibrometry methods prevent mosquitoes from beating their wings, so in order to simulate the effect of male flight on the male auditory organ, it is possible to combine the male’s flight sound-frequency with the female stimulus sound. For example, male *Cx. pipiens pipiens* JO sensitivity was improved by 7 dB with the addition of simulated flight sound at the main frequency optimum (18-22 °C) (Lapshin, 2012).

The results of electrophysiological and laser vibrometry studies can be difficult to compare against each other due to differences in methodologies used to assess threshold responses (e.g. determination of statistical definitions of neural thresholds and variations in the locations of electrodes). In addition, the main goal of these studies is not always about measuring absolute hearing thresholds, and as a consequence the number of replicates can be too few to analyse statistically.

Behavioural methods also face similar constraints, however, the assessment of physiological responses to sound stimuli offer a more natural context that enables more natural responses to sound. Behavioural responses provide more robust evidence of auditory outcomes because the whole auditory chain plus the motor responses are included. To our knowledge, there are only three published behavioural studies of mosquito sensitivity to sound intensity. First, Menda *et al.* (2019) measured the behavioural response of *Ae. aegypti* to 40 and 65 dB SVL by monitoring the take-off of resting mosquitoes in a cage located in the far-field of the sound-source. However, the behavioural methodology was not appropriate for the natural physiological context of swarming behaviour in this species; in the field both male and female *Ae. aegypti* fly continuously once the males detect the female’s flight tones (i.e., they rarely rest and take-off again). Indeed, male responsiveness to sound was found to be reduced when not flying (Lapshin, 2012).

Second, Feugère *et al.* (2021b) measured the flight and wingbeat frequency response of free-flying, swarming male *An. coluzzii* to a range of sound levels of a played-back group of females and found a response at 33±3 dB SPL. However, males may respond better to the sound of individual females rather than a group of females that would occupy a relatively wide range of wingbeat frequencies, as described for *Ae. aegypti* (Wishart and Riordan, 1959).

Third, Lapshin and Vorontsov (Lapshin and Vorontsov, 2021) showed an increase in flight speed in swarming male *Aedes communis* in response to the sound frequency of females in the field, with a hearing sound-level threshold of 26 dB SVL on average (26 dB SPL under far-field conditions; 12°C).

The aim of our study was to investigate the behavioural hearing threshold of *An. coluzzii* males; Pennetier et al. (2010) measurements suggest that their JO may be as sensitive as 10 dB SVL (range of 5-12 dB SVL, n=4 measurements on 2 males, criterion = 1.4 recording noise floor). As suggested 70 years ago by Roth (1948), male hearing may be enhanced during swarming behaviour (i.e., flying in loops over a floor marker, station-keeping while they wait for females to join the swarm) when male sensitivity to the sound of flying females is expected to be maximised. Therefore, we used a modified approach of Lapshin *et al.* who worked in the field with *Ae. communis* (Lapshin and Vorontsov, 2021). Our study was performed under the following conditions: 1) in a laboratory sound-proof chamber, with controlled measurement of sound levels; 2) with a range of type of sounds to be exposed to males; 3) by monitoring both the male flight-tone and the flight-dynamic quantitatively; and 4) with *An. coluzzii*, a swarming species belonging to the *Anopheles gambiae* complex. ‘Sound-level values’ depend on how sound level is defined and on the type of sound stimuli, therefore, a meaningful sound-level definition should be related to the sound-frequency band and temporal patterns which mosquitoes are sensitive to. For this reason, our main aim of quantifying hearing threshold was inter-connected with the following questions:

○ Is the second harmonic of female flight tones necessary to stimulate a response in males? We need this information to establish the frequency band(s) for which the sound level is defined to be appropriate to mosquito hearing.
○ Is temporal variation in natural female sound required for males to detect females or is a single-frequency at a constant amplitude sufficient?
○ Do the flight tones of a group of females have the same effect on male hearing as those of a single female, over a range of sound levels? The main interest in the last two questions is to investigate whether we can use single-frequency sounds to mimic female sound, which will make the hearing threshold easier to estimate in future studies.

## Materials and Methods

### Mosquitoes

All experiments were performed with virgin *An. coluzzii* Coetzee & Wilkerson. The colony was established at the Natural Resources Institute (NRI), University of Greenwich (UK) from eggs provided by the Institut de Recherche en Sciences de la Santé (IRSS), Burkina Faso. Eggs were obtained from a colony established in 2017 from wild gravid females collected from inhabited human dwellings in Bama, Burkina Faso (11°23’14"N, 4°24’42’W). Females were identified to species level by PCR (Fanello et al., 2002). The NRI colonies were kept in environmentally controlled laboratory rooms with a 12h:12h light:dark cycle (lights went off at 15h00), >60% relative humidity and ~24-26°C. Larvae were fed Tetramin^®^ fish-flakes and rice powder. Adult males and females were separated < 12h post-emergence to ensure all females were virgin and fed a solution of 10% sucrose and 1%-saline *ad libitum*. Adult mosquitoes were kept in cube cages of ~30 cm sides, populated with a) ~300 virgin females and b) ~20 males.

### Experimental setup

The basic experimental setup (Fig. 1) is the same as for a previous study with *An. coluzzii* (Feugère et al., 2021b) as described below.

**Fig. 1.**
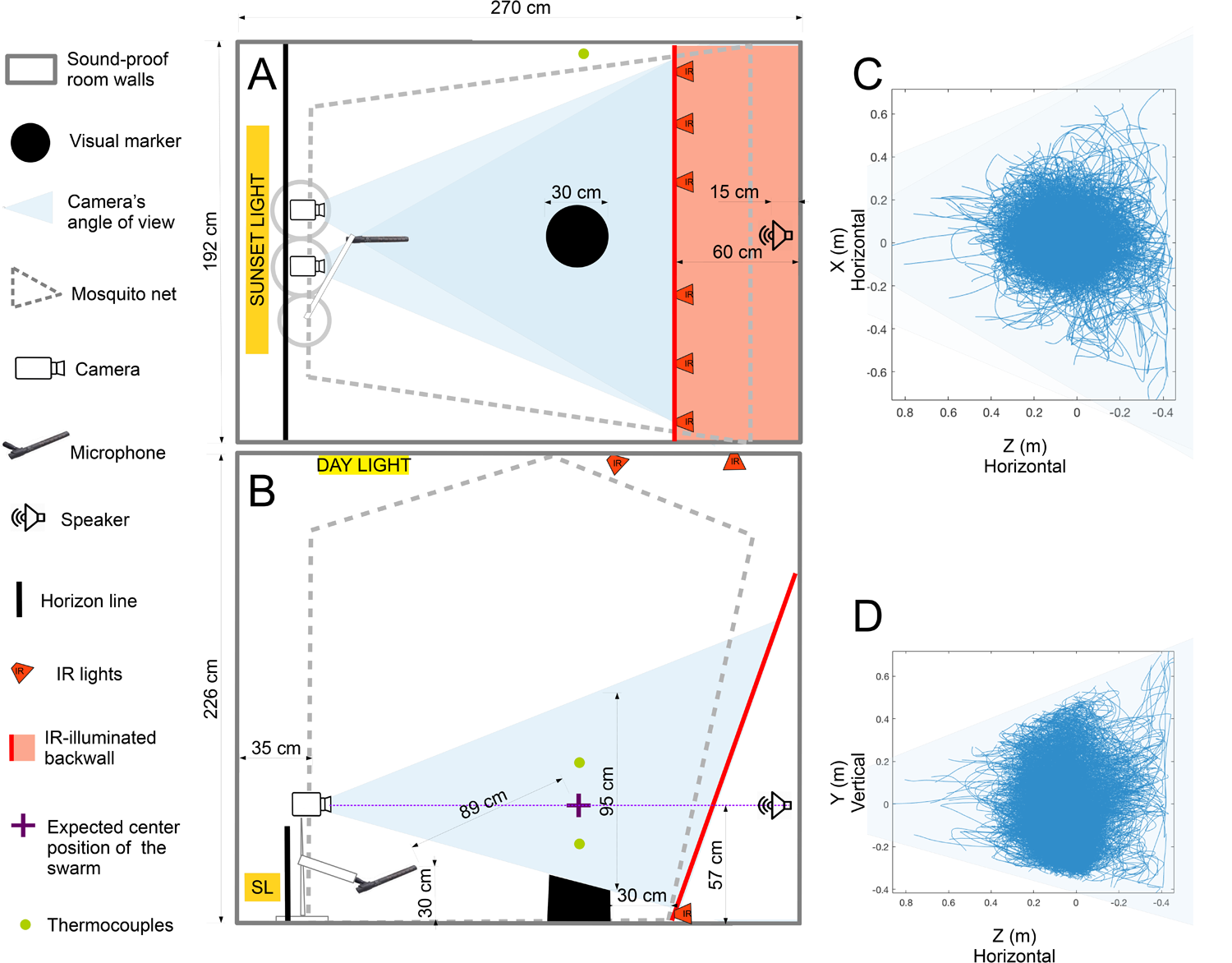
Sound-proof chamber setup for recording sound and video of *An. coluzzii* behaviour (modified version from (Feugère et al., 2021b)). (A) Bird’s-eye and (B) side views of sound-proof chamber. Blue shaded areas indicate the 3D fields-of-view of cameras recording mosquito flight paths. Two IR-sensitive cameras fitted with IR pass filters recorded flying mosquitoes as black silhouettes against evenly lit IR-background. A separate lighting system provided gradual semi-natural dusk visible to mosquitoes, consisting of dispersed dim white lights on ceiling and ‘sunset’ lighting below horizon (opaque wall ~40 cm tall). A microphone recorded flight sounds of mosquitoes swarming directly above black swarm marker. A thermocouple (85 cm above ground level) recorded temperature at ~ mean swarm height. A speaker located behind IR-illuminated thin-cotton sheet, outside net enclosure played back sound stimuli. (C) Bird’s-eye and (D) side views of the superimposed flight tracks of the entire dataset.

#### Sound-proof chamber

All experiments were conducted in a sound-proof chamber to limit interference from external sounds. The chamber consisted of double-skin sound-proof walls, ceiling and floor (*L* × *W* × *H* = 2.7 m × 1.9 m × 2.3 m), producing a reverberation time ≤ 0.07 s for frequencies above 200 Hz (IAC Acoustics, manufacturers). The SPL in the sound-proof room without any playback was always quieter than that with playback of the sound stimuli in the third-octave frequency band of the sound stimulus (Fig. S1 A). Below 176 Hz (upper limit of the 125 Hz octave band), the ambient noise level rase (Fig. S1 B; 25 dB at 125 Hz), due to low-frequency vibration of the building’s aeration system, which may have been detected by the *An. coluzzii* auditory system (Pennetier et al., 2010) as a low-frequency background noise to the sound stimulus.

#### Swarming arena

The swarming arena in the sound-proof chamber was designed to include the key environmental conditions and sensory cues known to control mating and swarming flight in the field. A large mosquito bed-net enclosure (NATURO, *L* × *W* × *H* = 1.8 m × 1.7 m × 2 m) filling most of a sound-proof chamber (Fig. 1) enabled mosquitoes to fly freely in a volume 100 times greater than that covered by the typical swarming space. Lighting was provided by an artificial-sunlight system to imitate natural daylight, sunrise and sunset (LEDs 5630, HMCO FLEXIBLE dimmer, and PLeD software, custom-built). Dimming the ambient light level at the appropriate circadian time elicits mosquitoes to take-off, followed by swarming behaviour in response to the presence of a visually conspicuous matt-black marker on the floor; both males and virgin females fly in loops above the marker, but this is rarely observed if males are present because males mate with females quickly and mated females cease swarming behaviour (Poda et al., 2019; Gibson, 1985). We used virgin female swarming behaviour to record their flight sound within a relatively limited distance from the marker.

#### Sound recording and monitoring

The wingbeats (aka, ‘flight tones’) of mosquitoes in the laboratory were recorded with a weatherproof microphone (Sennheiser MKH60; RF-condenser; super-cardioid polar pattern at 0.5-1 kHz, with amplitude decrease of > 15 dB beyond 90° from the microphone head; sensitivity at 1 kHz: 40 mV/Pa; A-weighting equivalent noise level: 8 dB) directed toward the swarm location. The microphone was located at a distance of 0.89 m from the centre of the swarm area for the experimental male mosquitoes and the sound recording of the 30-female swarm stimulus (Fig. 1), except for the recording of the 1-female sound-stimulus for which the microphone was located at 0.75 m from the centre of the swarm area. The microphone was plugged into a Scarlett 18i8 audio interface on a Windows7 computer running Pro Tools First 12.8 (Avid Technology, Inc).

#### Flight track recording

The 3D flight trajectories of male mosquitoes were recorded at a sampling rate of 50 Hz with Trackit software (SciTrackS GmbH, Switzerland (Fry et al., 2004)). Two video cameras (Basler, ace A640-120gm) were fitted with wide-angle lenses (Computar, T3Z3510CS, 1/3’’ 3.5-10.5mm f1.0 Varifocal, Manual Iris) to maximize 3D volume of video-tracking. IR lights (Raytec RM25-F-120 RAYMAX 25 FUSION) enabled the tracking system to detect flying mosquitoes as silhouettes against an IR-illuminated white back-wall made of thin cotton cloth (Fig. 1). The 3D-flight trajectories were smoothed using a cubic spline interpolation at a sampling frequency of 200 Hz on Matlab (version R2017a)

#### Temperature monitoring

Temperature was monitored by type-T thermocouples (IEC 584 Class 1, Omega) associated with a temperature logger (HH506RA, Omega) totalling a measurement accuracy error of ±0.9°C. The chosen thermocouple was located on a room wall at a height of 85 cm from the floor. The four recordings of the reference sound stimuli (two species, two sexes) were recorded at 28.0°C. The mean temperature and standard deviation of the behavioural assays were 28.0±0.3°C.

### Sound stimuli

#### Recording context

Two recordings of the natural flight-sounds of 3-6 days-old swarming females were recorded and used to produce the played-back stimuli for the behavioural assays. These sound recordings consisted of 1) a single swarming female or 2) a group of 30 swarming females; in both cases mosquitoes were released into the swarming arena 2 days before the experiment to acclimatize. The standard environmental conditions in the room were: 12h:12h light:dark cycle with a 1h artificial dawn/dusk transition in light intensity and ~60-75% RH.

#### Signal generation

We generated 4 types of stimulation signals (‘2-harmonic 1-female’, ‘2-harmonic 30-female’, ‘1-harmonic 1-female’ and ‘1-harmonic constant’) (Audio S1 to S4; signal spectrum in Fig. 2) over a range of sound levels, producing 10 stimuli in total. First, we selected the first 7s section of the sound of a single female swarming over the marker (Audio S5). Second, a 7s section of the sound of 30 swarming females was selected (Audio S6), ~10 min after the first female started to swarm. Four sound levels for each of the 1- and 30-female sounds were selected (10-45 dB SPL, Table 1), based on results of preliminary experiments. These 8 stimuli contained the two first harmonics. A high-pass filter was added to all the stimuli to remove the electrical noise below the first harmonic (at the noise level, see Fig. 2 and Table S1). In addition, we generated a 33 dB SPL stimulus, which has been shown in preliminary experiments to be the lowest level sound stimulus that females detect in the sound-proof chamber (but see Method section ‘Corrected SPLs for estimating the hearing threshold’ below). This sound stimulus included only the first harmonic because it has been shown electrophysiologically that the male auditory organ is more sensitive to the first harmonic than higher harmonics (Pennetier et al., 2010; Warren et al., 2009). Finally, we generated a synthetic 1-harmonic sound, called ‘1-harmonic constant stimulus’, with constant frequency and amplitude over time (set at the same mean peak-amplitude and mean frequency as the ‘1-harmonic 1-female’ sound). A gradual increase/decrease over 1 s in the level of the start and end sounds were added to avoid creating sound artefacts due to the signal truncation, and to make the stimulus more natural (possibly important for active antennal amplification (Jackson and Robert, 2006)). The 10 stimuli were played sequentially, with a 10 s interval of silence to be played-back during the behavioural assays. To avoid an effect of the order in which stimuli were played, 10 different sequences were generated, each containing the 10 sounds in random order. All stimuli were sampled at 8 kHz / 24 bits and designed in Matlab (R2017a, The Mathworks Inc, Natick, USA). Fig. 2 gives the sound spectrum and amplitude along time of each type of stimulus. Table S1 gives the filter/frequency parameters used to generate the stimuli. Table 1 gives the sound levels for each of them. Audio S5 and Audio S6 are the original 1-female and 30-female sound recordings, respectively. Audio S1 to S4 are the 4 types of stimuli; 2-harmonic 1-female, 2-harmonic 30-female, 1-harmonic 1-female, 1-harmonic constant, respectively.

**Fig. 2.**
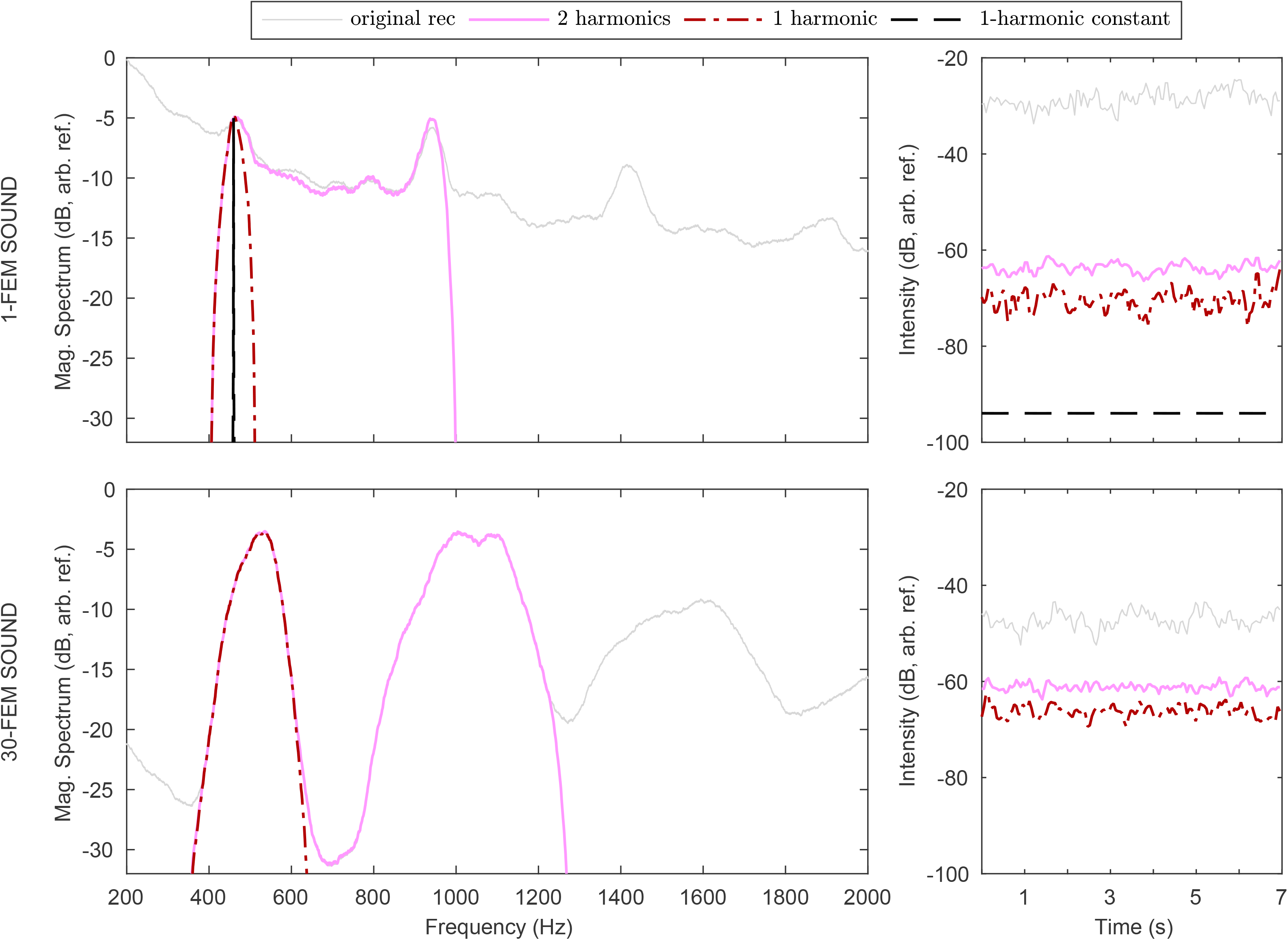
Spectral and temporal properties of sound stimuli. Spectral (first column) and temporal (second column) properties of sound stimuli of one single swarming *An. coluzzii* female (top row) compared to that of 30 females (bottom row). The originally recorded sounds are represented with a dotted line (Audio S1 and Audio S2 for a unfiltered 1-female and 30-female, respectively; not used directly as sound stimuli). The 1-harmonic 1-female sound is shown as a semi-dotted-dashed red line, while the 2-harmonic sounds are represented by a solid pink line. Magnitude spectra were calculated over 7 s and averaged over 50-Hz windows. The root-mean-square pressure levels were computed over a 0.1 s time window with 0.05 s overlap, along the 7 s duration of the stimuli. See Table S1 for characteristics of filters applied to Audio S1 and Audio S2 to generate the 1-harmonic and 2-harmonic stimuli.

**Table 1.** Description of stimulus sound-levels. This table gives the sound pressure levels (SPL ref 20 μPa) and associated errors of all played-back sound stimuli at the male’s mean location in the frequency range of the female’s first harmonic. See Methods ‘Corrected SPLs for estimating the hearing threshold’ for the corrected SPL mean value and Methods section ‘Estimate of SPL errors at mosquito’s location’ for last two columns. SPLs are equal to SVLs in our setup (see Method section ‘Monitoring SVL from SPL measurements). For frequency characteristics, see Fig. 2 and Table S1.

| Subset | Single/ Group | Number of harmonics | Recording/ Synthetic | Sound level (dB SPL) |  |  |  |
| --- | --- | --- | --- | --- | --- | --- | --- |
|  |  |  |  | SPL measurement of the two 1/3-octave bands closest to the first-harmonic at fixed distances from the speaker (0.9 m) |  |  | Error due to ±0.2 m oscillation (and total error) |
|  |  |  |  | Mean value | Corrected mean value | Error over the 7 s |  |
| NA | Silence playback | NA |  | 6.9 |  | ±0.3 | ±0.3 |
| A | Single | 1 | Recording | 25.0 |  | ±0.9 | ±2 |
|  |  | 2 |  | 32.6 |  | ±0.3 | ±2 |
| B | Single | 1 | Recording | 25.0 |  | ±0.9 | ±2 |
|  |  |  | Synthetic | 26.0 |  | ±0.2 | ±2 |
| C | Single | 2 | Recording | 10.6 | NA | ±0.5 | ±2 |
|  |  |  |  | 22.4 | 14.4 | ±0.4 | ±2 |
|  |  |  |  | 32.6 | 24.6 | ±0.3 | ±2 |
|  |  |  |  | 44.2 | 36.2 | ±0.6 | ±2 |
|  | Group (~30) |  |  | 17.1 |  | ±0.5 | ±2 |
|  |  |  |  | 23.1 |  | ±0.4 | ±2 |
|  |  |  |  | 32.9 |  | ±0.5 | ±2 |
|  |  |  |  | 44.9 |  | ±0.5 | ±2 |

#### Sound diffusion

Sequences of sound stimuli were played-back from a speaker (Genelec 8010A) plugged into a Scarlett 18i8 sound card running pro-Tools First and Audacity on Windows 7. The speaker is composed of two membranes (ø 76 mm and 19 mm). The centre of the larger speaker’s membrane was located 57 cm above the floor, 15 cm from the back wall and 0.9 m from the swarming centre (Fig. 1). The speaker’s self-generated noise was less than 5 dB SPL (A-weighted) and the sound card’s Equivalent Input Noise was −127 dBu.

#### Data Subsets

While stimuli were played-back in random order during a single experiment, they can be grouped into three overlapping subsets (Fig. 3), each of which corresponds to one of the questions presented at the end of the Introduction; Subset A: study of the effect of the second harmonic on male hearing (1-harmonic *vs* 2-harmonic stimuli), Subset B: investigation of the effect of ‘types of sound stimulus’ (single-frequency *vs* pre-recorded played-back stimuli) and Subset C: effect of the number of females (1 *vs* 30) in the recorded-sound stimuli and of the sound levels of the sound stimuli on male hearing to estimate the hearing threshold.

**Fig. 3.**
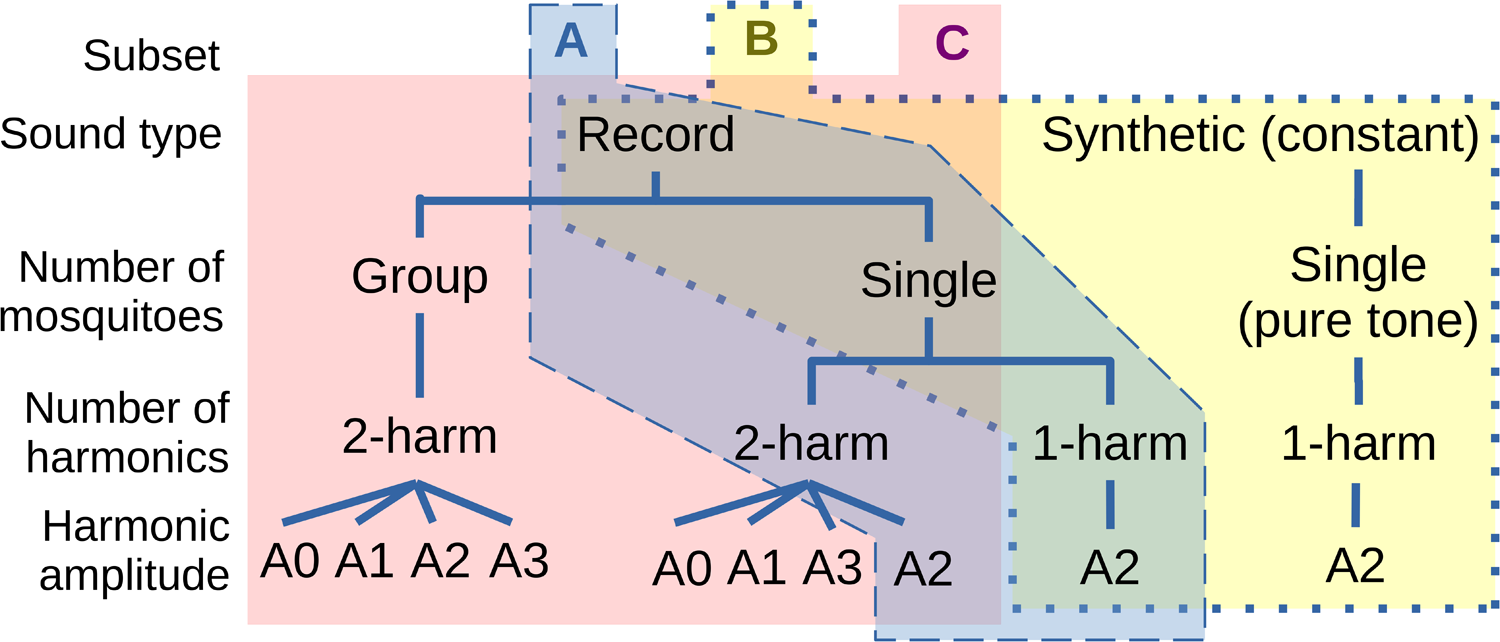
Data subsets for our analysis (A, B, C). Subset A was used to study the effect of the number of harmonics in the sound stimuli. Subset B was used to compare the sound type (playback of female sound or constant sound of the same wingbeat frequency). Subset C was used to study the effect of number of mosquito(es) (1 female or 30 females).

### Behavioural assays

To investigate the sensitivity of swarming males to female sounds, we played-back the female sound stimuli to swarming males in the sound-proof chamber. About twenty 3-4 days-old males were released the day prior to experiments at ~ 18h00 in the sound recording flight arena. At 15h00, after the ceiling lights had dimmed to the lowest intensity, the horizon light completed a 10 min dimming period and then kept at a constant dim light intensity until the experiment was finished. When at least one male started to swarm robustly over the marker, the first sequence of all 10 sound stimuli (i.e. the 4 types of stimuli, with 4 sound levels for 2 of them, see Method section ‘Signal generation’) was played-back from the speaker (see Movie S1 with a male exposed to one sound stimulus; see Fig. S2 for examples of responses for each type of stimulus). After 10 stimuli were played and if the male(s) was still swarming, or as soon as at least one male started swarming, a new sequence of 10 stimuli was immediately played and so on, until up to 10 sequences were played or after 50 min of constant horizon light, either of which marking the end of the experiment for the day (= 1 replicate). Males were then collected and removed from the flight arena. A new group of ~20 male mosquitoes were released in the sound-proof chamber, to be used for a new replicate the next day (one replicate per day, for 10 days in August-September 2018).

### Sound pressure level (SPL)

#### Measurement

Stimulus SPLs were measured at the mean male swarming position with a sound meter (Casella, CEL633C1, Class 1) set as follows: reference pressure of 20 μPa; no octave weighting (i.e., dB Z); slow octave time-constant (IEC 61672-1: 2002); octave and third-octave bands; calibrated twice a day (CEL-120/1, Class 1, at 94 dB / 1 kHz) before and after each measurement. The speaker and the software/soundcard gains were set to be the same as during the behavioural experiment.

#### Third-octave bands

All SPLs reported in this study included only the frequency bands that are audible to male mosquitoes, i.e., mostly the first harmonic of the female (Warren et al., 2009; Pennetier et al., 2010). They were calculated as follows: 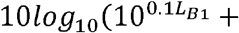 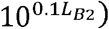 where L_B1_ and L_B2_ are SPL measurements in frequency bands *B1* and *B2*; *B1*=500 Hz and *B2*=630 Hz are the third-octave bands nearest the female’s wingbeat frequency of the first harmonic (Table 1; and Fig. S1 for all third-octave values).

#### Corrected SPLs for estimating the hearing threshold

The sound of 1-female was recorded at a distance of 0.7±0.2 m, which gave a relatively low signal-to-noise ratio compared to the high signal-to-noise ratio of the sound of 30-females recorded at 0.9±0.2 m. As explained in the Method section ‘Sound stimuli’, noise was removed below the first harmonic and above the second harmonic but not in-between to limit artefacts in the sound stimulus. SPL was computed over the frequency-band of the first harmonics, which, for the 2-harmonic 1-female sound, included a part of the noise between the first and second harmonics. Results from Subset A indicated that males did not need this noise to respond to sound because they reacted to the 2-harmonic 1-female sound as much as to the 1-harmonic 1-female sound. Since these two stimuli had the same first-harmonic amplitude but a SPL difference of 8 dB (Table 1), and because SPL was defined over a frequency band below the second harmonic, we established that the noise between the first and second harmonics is responsible for 8 dB in our SPL measurements. In order to estimate an accurate hearing threshold, we applied a correction of 8 dB to the sound level of the 2-harmonic 1-female stimuli (Subset C). All sound levels, with correction or not, are summarized in Table 1.

#### Control of distance between live mosquito and playback speaker

Swarming mosquitoes confine themselves to a limited area of the flight arena naturally, which enables us to estimate the incident SPL at the mosquito’s location, because the distance between swarming mosquitoes and the sound stimulus source was limited to a known range. The speaker (Genelec 8010A) that reproduced the females’ flight tones was placed 0.9 m from the centre of the swarm marker. Their flight positions were recorded by 3D-tracking Trackit Software (Fry et al., 2004) (Figs 4 A, 4 B) which enabled us to determine the distance between a mosquito and the speaker emitting mosquito sound to be 0.9±0.2 m, 95%-CI (Fig. 4 C).

**Fig. 4.**
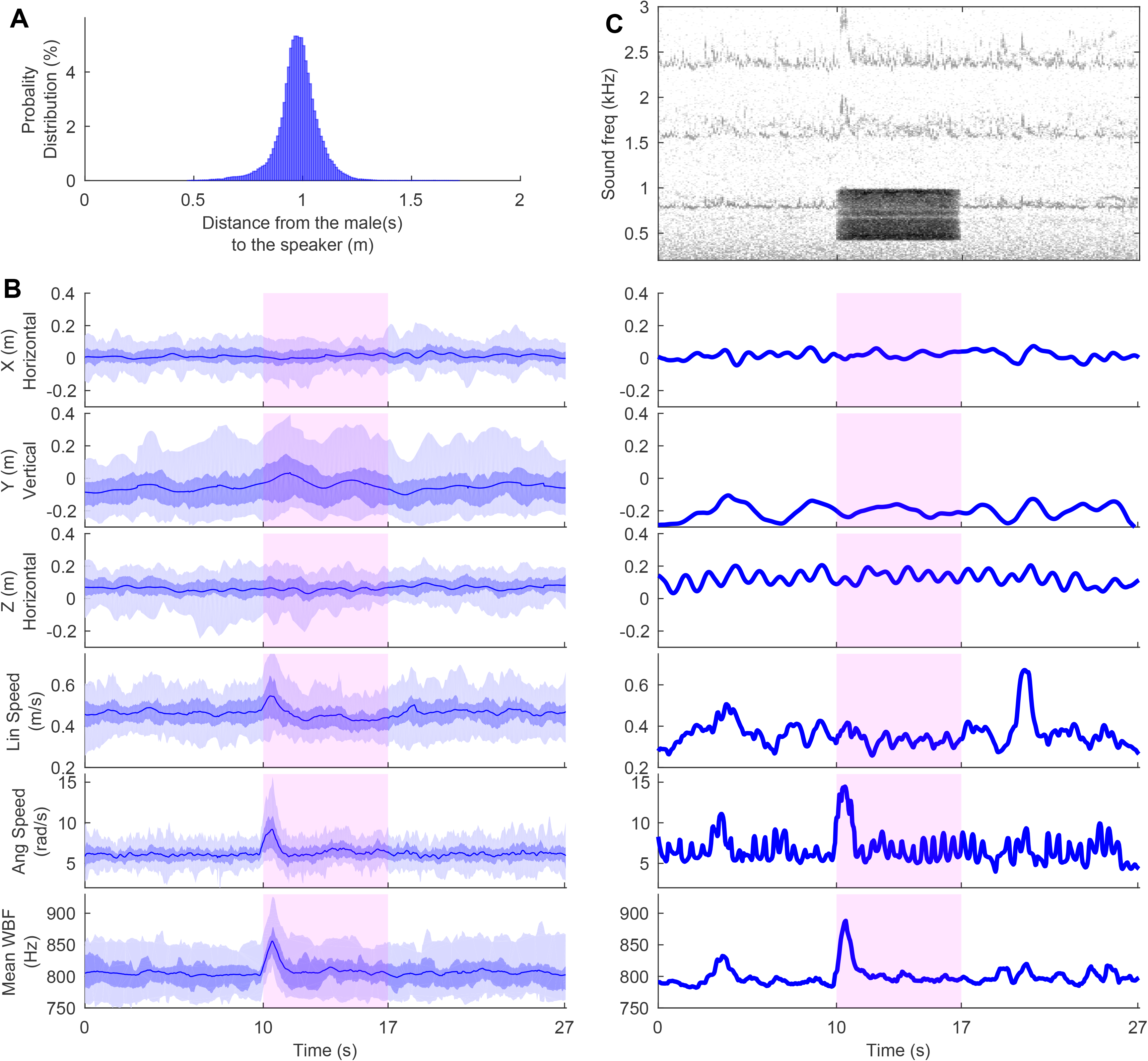
Flight and sound responses of *An. coluzzii* males to sound-stimuli. Male flight-characteristics and wingbeat-frequencies (blue) before, during and after playback of female (red rectangle) sound stimuli. **(A)** Example of male response to the loudest 2-harmonic 1-female sound-stimulus over 27 s of recording. Stimulus was played-back 10 s from beginning of flight recording and lasted 7 s (red rectangular shading). First five rows show flight parameters (relative X,Y and Z positions, plus linear and angular flight speeds). ‘Z’ dimension represents relative distance to the speaker (located 0.9 m from *Z*=0). Before-last row shows mean wingbeat frequency (WBF). Periodic flight pattern, typical of swarming behaviour, is evident in X, Y and Z plots. In the angular-speed and wingbeat frequency plots, the two red lines correspond to the upper-quartile over 1s and the arrows represent the differences between the two red lines, which are the parameters computed for monitoring the male response (see Methods section ‘Extraction of traits used to quantify male responses’). Last row shows the spectrogram of sound recordings before, during and after the sound stimulus; the colour gradient represents the sound level given a frequency and a time (the darker the colour, the louder the frequency). Movie S1 gives the associated raw image and sound recording. See Fig. S2 for examples of responses to the 4 types of sound stimulus. **(B)** Same as (A) but without spectrogram and for all-male responses to the loudest 2-harmonic 1-female sound-stimulus. Darkest coloured lines represent running median, darkest areas represent second and third quartiles and light areas represent the 90^th^ percentile of data. The sample size of the distribution of flight coordinates and velocities corresponds to the number of male flight tracks (n=104), and that of the WBF distribution corresponds to the number of swarms (n=61) where mean WBFs over the number of mosquitoes per swarm were calculated (1 to 6 males per swarm). Linear and angular speed, and wingbeat frequency clearly increased in response to the onset of this sound stimulus, plus there was a slight tendency to increase in flight height (Y (m)). **(C)** Probability distribution of distance between a male and the speaker during sound stimulus playback for all stimuli; distances ranged between 0.9±0.2 m. This distance interval was used to estimate the uncertainties of the acoustic prediction in Table 1. The sample size of the distribution of distances corresponds to the number of male flight tracks (n=104).

#### Estimate of SPL errors at mosquito’s location

Two types of SPL errors were taken into account. The first is related to the time variation of the sound stimulus levels which were between ±0.3 dB and ±0.9 dB (maximum error), depending on the stimulus (see Fig. 2 for an example of stimulus sound-level over time). The second type of measurement uncertainty arises when the sound level should be estimated from the mosquito’s position, and not from the fixed microphone position. Indeed, SPLs were measured at the expected centre of the station-keeping swarm-flight of the test male mosquitoes. However, the distance between the male and the speaker varied as 0.9±0.2 m (95%-CI, Fig. 4 C), due to the males’ swarming-flight pattern, which changed the sound level they were exposed to, accordingly. We evaluated this error by playing-back the *An. coluzzii* female sound stimulus and measured the sound level in a sphere around the expected swarming area centre: the maximum error was ±2 dB. This error is considered to be conservative (at least 95%-CI) and was used to interpret the results of the experiments (see Table 1).

#### Physical sound quantities produced by a speaker and sensed by mosquitoes

We monitored the sound level of the played-back stimuli by recording the sound pressure level (SPL), however, mosquito hearing organs are sensitive to particle velocity level (SVL) (Fletcher, 1978). The root-mean square value (RMS) particle velocity *v*_*RMS*_ and the RMS sound pressure *p*_*RMS*_ can be related as follows, assuming the speaker to be a point source radiating spherically a sound frequency *f* at a distance *r* from the source (air impedance *Z*_*air*_(28°C) = 408 N.s.m^−3^; sound speed *c*(28°C)=348 m/s) (Beranek and Mellow, 2012):

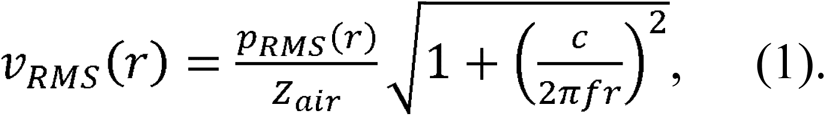

The SPL 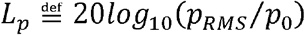 and the associated particle-velocity level *L*_*v*_ = 20*log*_10_(*v*_*RMS*_*Z*_*air*_/*p*_*0*_) (reference *p*_*0*_ = 2.0 10^−5^ Pa) can be calculated as follows:

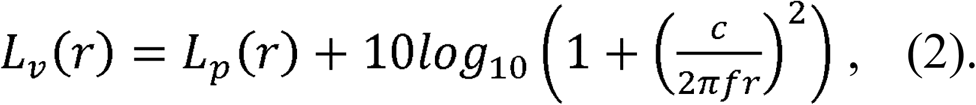

Considering that the female sound stimulus does not have any frequency components below *f* = 440 Hz (the smallest frequency value of the group of first harmonics of the swarming females at −12 dB below the peak at 536 Hz, Fig. 2), the SVL is equal to the SPL at 0.9 m away from a monopole sound source of these frequencies, under a negligible error of less than 0.1 dB (due to the mosquito oscillating distance of ±0.2 m to the speaker, calculated from equation (2)). As a consequence, and since mosquitoes are sensitive to SVL and for easier comparison with other studies, we report the SPL as SVL. Arthur et al. (2014) measured the particle-velocity attenuation rate in front or behind *Ae. aegypti* to be between a monopole and a dipole. Note that our monopole assumption for mosquito wing-flapping is conservative since higher orders (dipole, quadripole) produce sound levels that decrease more rapidly with distance (Bennet-Clark, 1998).

### Extraction of traits used to quantify male responses

Following the results of preliminary experiments, we used two components of male flight: 1) **angular-speed**, calculated from their 3D trajectories and 2) **wingbeat frequency**, extracted from sound recordings (see Fig. 4 B for example and statistics of wingbeat and flight dynamic characteristics before, during and after exposure to the loudest 1-female sound-stimuli (44±2 dB SVL)). The two components were synchronized using the same techniques as in a previously published study (Feugère et al., 2021c).

***Angular-speed*** refers to how much the mosquito flight direction changes per unit time. It was calculated from the linear-velocity components provided by the Trackit software as follows: *avel=Δθ* / *Δt*, where *Δt* = *t*_*n*_ − *t*_*n*+1_ is the duration between two consecutive time indexes *n* and *n*+1, and *Δθ* is the turn angle defined as:

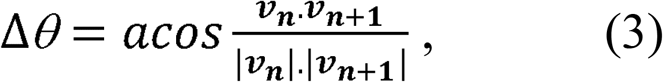

where ***v***_***n***_ is the three-dimensional linear velocity vector of the mosquito at time index n and |***v***_***n***_| is its magnitude. The criteria used to include a tracked flight in the data analysis were that the mosquito was swarming over the marker for at least 1 s before and after the sound stimulus onset.

#### Wingbeat frequency

Only the first and/or the second harmonic of female sound stimuli were played-back (~400-1200 Hz) in order to free the frequency domain of the male’s third harmonic from the female’s sound. This allowed us to capture the male’s third harmonic without overlapping with the sound stimulus (example of spectrogram in Fig. 4 A). The peak of the third harmonic was detected every 40 ms between 2190 and 2920 Hz using the Fast Fourier Transform algorithm (256-ms FFT-window, Hanning-windowed). When several mosquitoes (from 1 to 6) were present over the swarming marker, the detected value was the peak of the energy in the frequency band 2190-2920 Hz and not the mean of the peak from individual mosquitoes (because it was not possible to track the wingbeat frequencies of individual mosquitoes). Then, the male’s third-harmonics (i.e., 3 × wingbeat frequency) were divided by 3 to get the wingbeat frequency (i.e., the first-harmonic frequency). Finally, a 3-point median filter was applied over time to reduce wingbeat tracking error. Fig. 4 A gives an example of detected wingbeat frequencies of males while Fig. 4 B shows the distribution of the detected wingbeat frequency over time for all recordings.

#### Upper-quartile difference

Since preliminary experiments suggested that mosquitoes responded to sound by increasing their wingbeat frequency and their angular speed somewhere during the first second of the sound stimuli, the upper-quartile angular-speeds and the upper-quartile wingbeat frequencies were automatically detected during the first 1s stimulus time interval. Indeed, ‘upper-quartile’ is 1) a more robust metric than median or mean to measure the amplitude of a short peak, which the onset time cannot be predictable precisely and 2) a more reliable metric than ‘maximum’ to avoid false detection. Then, this value was subtracted from the upper-quartile value computed during the 1s segment just before the stimulus onset, for each individual recording to reduce noise related to individual mosquito variability (Fig. 4 A shows graphically how the parameters were computed).

### Statistics

Wingbeat-frequency and angular-speed values for a given stimulus were averaged over the different responses of the same day to form a replicate. The wingbeat and angular-speed response-parameters were analysed using a Bayesian Linear Mixed-Effects Model (*blmer* function, *lme4* package, R). Stimulus sound levels (continuous), number of females in the recording (1 or 30), number of harmonics (1 or 2) and sound type (recording or synthesis) and their interaction were considered as fixed effects. Days, for which replicates were performed, were considered random effects. The dataset was split into the 3 subsets A, B, C, as shown in Fig. 3. A total of 6 models were built (2 parameters × 3 subsets). Stepwise removal of terms was used for model selection, followed by likelihood ratio tests. Term removals that significantly reduced explanatory power (*p*<0.05) were retained in the minimal adequate model (Crawley, 2007). No data transformation was needed to ensure variance homogeneity of variables (Fligner-Killeen test, *Fligner.test* function, R) and normality of model residuals (Shapiro-Wilk test, *shapiro.test* function, R), except for Subset C wingbeat-frequency which was transformed via optimality (*MLE_LambertW* function, *LambertW* package, R (Goerg, 2016)); see Fig. S3 for normality *qqplots* and Table S2 for normality and variance homogeneity test results.

For subsets A and B, an additional one-sample t-test (with BF-correction for multiple comparisons) was performed independently for each distribution to measure the significance of the mean to 0, which is the “no response” reference. For subset C, the quietest 2-harmonic 30-female sound-stimulus was not included in the model because its sound level was too close to the background noise level to be corrected as the three other 2-harmonic 30-female sound-stimuli. The hearing threshold was estimated by the crossing of the *y*=0 axis (i.e., no response, including with the LambertW transformation) with the prediction of the fixed-effect components of the mean and associated 95%-CI (*bootMer* function with *nsim*=500*, lme4* package, R). The Lambert transformation does not change the 0 value of the distribution. All analyses were performed using R (version 3.5.3).

Model Subsets resulted in a sampling size of *n=*10 for Subset A and B and *n*=9 or *n*=10 for Subset C (see legend of Fig. 5 for details; see Method Section ‘Behavioural assays’ for how a replicate was defined).

**Fig. 5.**
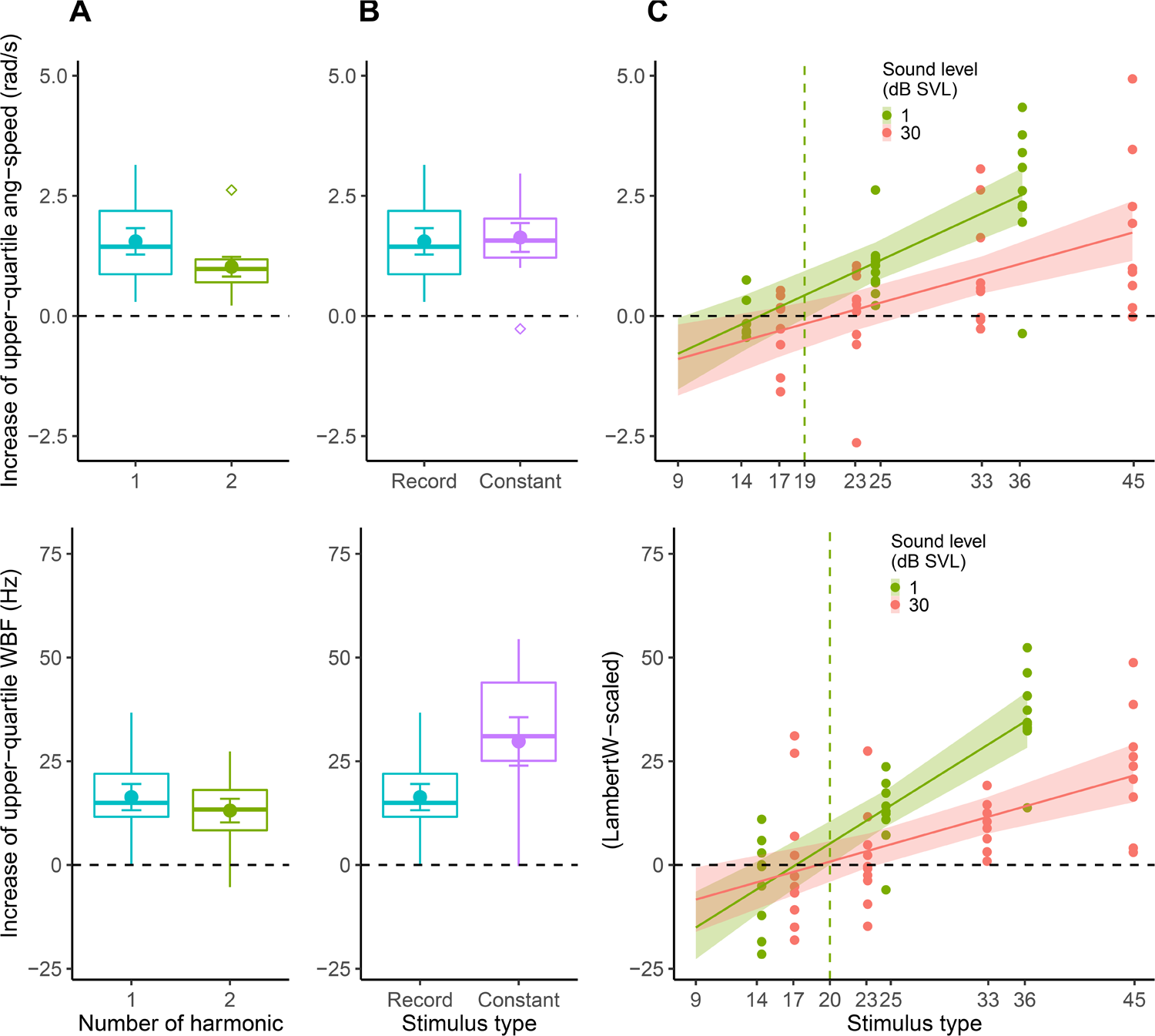
Results of behavioural experiment. Top and bottom rows show the increase in upper-quartile angular-speed and wingbeat frequency, respectively, when playing-back a given sound stimulus. Black dotted lines represent the absence of change in parameters before and during the stimuli. Each sample is the average of several measurements on the same day. Each sample corresponds to a different group of mosquitoes (consisting of 1 to 6 in each sample). See Method Section ‘Statistics’ and Results Sections for statistical tests. **(A)** Male *An. coluzzii* responses to 1- or 2-harmonic sounds of a single female (data subset A, n=10 in each boxplot). Boxplots of the parameters show the median, 2^nd^ and 3^rd^ quartiles. Outliers shown as diamond shapes are outside the interval [Q1 – 1.5 * IQD, Q3 +1.5 * IQD] which is represented by whiskers (Q1 = first quartile; Q3 = third quartile and IQD = interquartile distance). Disk and error bars in each distribution show mean and standard error. **(B)** Male *An. coluzzii* responses to 1-harmonic 1-female sound or to single-frequency sound (data subset B, n=10 in each boxplot). Boxplots, disk and error bars have the same meaning as in (A). **(C)** Male *An. coluzzii* responses to 2-harmonic sounds of single or 30 females along SVLs (data subset C, n=9 for the quietest 1-female stimulus and the two loudest 30-female stimuli, n=10 for other stimuli). Continuous lines and associated coloured areas represent the mean and 95%-CI. SVL were corrected as explained in Method section ‘Corrected SPLs for estimating the hearing threshold’. The green dotted lines represent the lowest estimate of the hearing threshold from the response to 1-female 2-harmonic sound-stimuli.

## Results

### Males mostly use the female’s first harmonic to hear her flight tone (Subset A)

Subset-A sound-stimuli with one or two harmonics were heard by males as the response distributions are different from the null distribution (Fig. 5 A), for both angular-speed (upper-quartile angular-speed difference: one-sample *t*=5.7, *df*=9, BH-corrected *p*<0.001, mean=1.5 rad/s; one-sample *t*=5.0, *df*=9, BH-corrected *p*<0.001, mean=1.0 rad/s, respectively) and wingbeat frequency (upper-quartile wingbeat-frequency difference: one-sample *t*=5.2, *df*=9, BH-corrected *p*<0.001, mean=16 Hz; one-sample *t*=4.6, *df*=9, BH-corrected *p=*0.0013, mean=13 Hz, respectively).

Our results show no differences in response of males exposed to the first-harmonic’s sound of a female flight tone or the combination of the first and second harmonic sounds with noise in-between of the same female flight tone (upper-quartile angular-speed difference: LRT, *χ*^2^=2.6, *df*=1, *p=*0.11; upper-quartile wingbeat-frequency difference: LRT, *χ*^2^=1.1, *df*=1, *p=*0.29).

### Males react to a ‘pure-sound’ (1-harmonic constant sound) at least as much as to a ‘natural sound’ (1-harmonic 1-female sound) (Subset B)

Subset-B stimuli, i.e. 1-harmonic 1-female sound and 1-harmonic constant sound, were both heard by the males because the response distributions were different from the null distribution (Fig. 5 B), for both the angular-speed (upper-quartile angular-speed difference: one-sample *t*=5.7, *df*=9, BH-corrected *p*<0.001, mean=1.5 rad/s; one-sample *t*=5.4, *df*=38, BH-corrected *p*<0.001, mean=1.6 rad/s, respectively) and the wingbeat frequency (upper-quartile wingbeat-frequency difference: one-sample *t*=5.2, *df*=9, BH-corrected *p*<0.001, mean=16 Hz; one-sample *t*=5.1, *df*=38, BH-corrected *p*<0.001, mean=30 Hz, respectively).

Our results show there is little difference in the male response between the 1-harmonic 1-female sound-stimulus and the 1-harmonic constant sound of the same mean frequency/SVL. While males change their angular-speed with the same amplitude in response to these two stimuli, they change their wingbeat frequency two times more with the 1-harmonic constant sound (upper-quartile angular-speed difference: LRT *χ*^2^=0.052, *df*=1, *p=*0.82; upper-quartile wingbeat-frequency difference: LRT *χ*^2^=4.5, *df*=1, *p=*0.033, respectively).

### Males react to the 1-female sound more than to the 30-female sound, with a hearing threshold less than 20 dB SVL (Subset C)

Using data subset C (see Table 1 for sound levels), our results (Fig. 5 C) show that free-flying males respond to the sound stimuli, providing the sound level was high enough, by increasing both their angular-speed and their wingbeat frequency as the tested sound levels increased (upper-quartile angular-speed difference: LRT *χ*^2^=36.8, *df*=1, *p*<0.001, effect size=0.12 rad/s per dB SVL; and LambertW-transformed upper-quartile wingbeat-frequency difference: LRT *χ*^2^=23.8, *df*=1, *p*<0.001, respectively). The number of females had small effect, but this was not interpretable, because of distinct values of sound levels for each number of females (upper-quartile angular-speed difference: LRT *χ*^2^=3.3, *df*=1, *p*<0.001, 1.2 rad/s for 1-female *vs* 0.6 rad/s for 30-female stimuli; and LambertW-transformed upper-quartile wingbeat-frequency difference: LRT *χ*^2^=3.2, *df*=1, *p*=0.073, 20 Hz for 1-female *vs* 10 Hz for 30-female stimuli, respectively). However, globally, the males responded more to the 1-female sound than to the 30-female sound as the sound level increased (i.e. interaction between the sound level and the number of female; upper-quartile angular-speed difference: LRT *χ*^2^=3.3, *df*=1, *p*=0.070, effect size = additional 0.05 rad/s per dB SVL for 1-female sound-stimulus; and LambertW-transformed upper-quartile wingbeat-frequency difference: LRT *χ*^2^=10.3, *df*=1, *p*=0.0013, respectively).

For 2-harmonic 30-female sound-stimuli (Fig. 5 C, red colour), the mean sound-level threshold was 21 dB SVL with a 13-27 dB SVL 95%-CI, if considering the angular-speed as response parameter. Using the wingbeat frequency parameter, the mean sound-level threshold was 19 dB SVL with a 9-23 dB SVL 95%-CI. For 2-harmonic 1-female sound-stimuli (Fig. 5 C, green colour), the mean sound-level threshold was 15 dB SVL with an 9-19 dB SVL 95%-CI, if considering the angular-speed to be a response parameter. Using the wingbeat frequency parameter, the mean sound-level threshold was 17 dB SVL with a 13-20 dB SVL 95%-CI. Considering these latter stimuli, which are the most ecological ones, a conservative estimate of the hearing threshold is then 20 dB SVL.

## Discussion

### Behavioural assessment of hearing threshold in swarming mosquitoes

Inter-mosquito acoustic communication is believed to occur at short range only (Feugère et al., 2021b), during mating behaviour when mosquitoes are flying in loops near a visual marker. *Anopheles coluzzii* males gather in tens to thousands over station-keeping swarm sites, while virgin females join the swarm in much fewer numbers as they mate only once in a life-time. Once a male detects a female’s presence from her wing-flapping sound, the male starts to chase the female (Pantoja-Sanchez *et al*., 2019). Thus, there is strong competition between males to detect relatively rare females (~1% male:female ratio (Kaindoa et al., 2017; Charlwood and Jones, 1980). Accordingly, acute hearing sensitivity is highly advantageous to males, along with other factors such as their own wingbeat acoustic power (Lapshin, 2012) and frequency (Somers et al., 2021) in the context of distortion-product hearing.

Under laboratory conditions (27-29°C), we show that male *An. coluzzii* respond strongly to 1-harmonic constant sound of 26±2 dB SVL at the female’s mean wingbeat frequency (Fig. 5 B and Table 1) and we estimate the hearing threshold to be 20 dB SVL or less with a 95%-CI using 2-harmonic 1-female sounds (13-20 dB SVL). Researchers have used electrophysiological mosquito preparations to measure hearing thresholds in the Johnston’s organ, which does not involve free-flying, pre-mating behaviour, such as swarming (but see Feugère et al. (2021) and Lapshin and Vorontsov (2021)). This may explain why these electrophysiological studies usually found far higher sound thresholds than in our study (see Introduction section). Lower hearing thresholds measured by electrophysiological methods can partly be explained by the absence of flight tones in males, which is known to be important to enhance the sensitivity in males to female sound. This creates mixed-harmonics for which the JO is tuned to, as shown by electrophysiology mosquito preparations exposed to flight sound simulation, which lowers the hearing threshold by 7 dB in *Cx. pipiens pipiens* (Lapshin, 2012). However, this may not be the only explanation. Mosquitoes exhibit ‘active hearing’, which can be triggered only during specific physiological states (Göpfert and Robert, 2001; Su et al., 2018), one of which may be swarming. It may be that males can enhance hearing to detect a female that is approaching a male swarm before she is chased by a competitor.

The only other species to have been explored in relation to these aspects of swarming flight is *Ae*. *communis* (Lapshin and Vorontsov, 2021); in the field, the mean hearing-threshold of males at the female’s wingbeat frequency was shown to be particularly low, 26 dB SVL. However, their method consisted in monitoring flight-speed changes in natural swarms by eye, which may not have enabled them to measure the smallest response amplitudes, thereby over-estimating the threshold (Lapshin and Vorontsov, 2021). On the contrary, we measured both flight dynamics and wingbeat frequency from quantitative measurements. Also, ambient temperatures were very different (~12°C for Lapshin and Vorontsov (2021) *vs* 27-29°C for our recording), which can change hearing sensitivities.

Finally, electrophysiological measurements in the JO are usually averaged over JO scolopidia, however, this could misrepresent the effective signal that triggers a behavioural response. Indeed, in addition to individual sensitivity in frequency and threshold, JO scolopidia are sensitive to the direction of the sound wave, and then only the JO scolopidia which are aligned with the sound wave-front display a low response-threshold. As a consequence, averaging all JO-scolopidium thresholds may over-estimate hearing thresholds (Lapshin and Vorontsov, 2019).

### Male response to sound and the effect of number of females

Males change their wingbeat frequency with a greater amplitude when exposed to 1-female sound than to 30-female sound, however, the change in angular-speed was small and its statistical significance was marginal (Subset C). This occurs despite the relatively greater amount of noise in-between the 1^st^ and 2^nd^ harmonic in the sound stimulus of the 1-female; the difference may have been stronger if the prominence of the harmonics had similar values in the tested stimuli. Two comments merit emphasis; the first is that a group of frequencies that are attractive alone (e.g., grouped-female sounds) have a masking effect on mosquito auditory perception. These results support reports published 80 years ago with *Ae. aegypti* males; it was observed that these mosquitoes were not attracted to two or more sounds at a time, even though each of these sounds were attractive on their own (Wishart and Riordan, 1959).

Second, it is interesting that males respond more with their wingbeat frequency than with their flight trajectory or dynamics. The change in wingbeat-frequency is consistent with a current theory that during a chase between a male and a female, the male moves to the sound source by tracking the female’s wingbeat sound and adjusts his own wingbeat frequency to hear her better, through an auditory mechanism based on antennal distortion products (Warren et al., 2009, Simões et al., 2019). In our case, the sound wave-front is almost planar at the male’s position, due to the distance and membrane dimension of the speaker, contrary to the sound wave of a female of the same sound level which would be far more spherical. This may create contradictory signals in the mosquito auditory system, i.e., the sound level suggests that the female is very close, but the sound wave-shape gives poor information about her actual location.

### The question of hearing higher harmonics and the significance of background noise

Males are known to detect mainly the female’s first harmonic to hear her flight tone. Indeed, *Ae. aegypti* respond (with clasping and seizing movements in flight) to low frequencies under 500 Hz (i.e., 1-harmonic sounds) using tuning forks (Roth, 1948), while other species, such as *Cx. pipiens pipiens*, have a narrower frequency range of response (500-600 Hz) when swarming (Gibson, 1985). In *Toxorhynchites brevipalpis, Cx. pipiens pipiens,* and *An. gambiae s.l.*, electrophysiology revealed that male antennae are sensitive to a large frequency-band up to 2 kHz that encompasses the two first harmonics, however, the electrical tuning of their JO is very narrow and centred on the difference wingbeat frequency of the two sexes which is close to the female’s first harmonic (Gibson et al., 2010). With respect to behaviour, Wishart and Riordan (1959) trapped as many *Ae. aegypti* males with the sound of 1-harmonic tones as with the complete flight sound.

Moreover, when removing the first harmonic from female flight tone recordings, *Ae. aegypti* males did not respond anymore, but the authors reported their results without any further information. This absence of a male’s response if the female’s first-harmonic is removed from the stimulus is similar to our results with *An. gambiae*, which shows a similar male response if the second harmonic of the female flight-tone is removed. On the contrary, it has been reported that male *Ae. aegypti* can hear the female second-harmonic, but without inferential statistics (Cator et al., 2009), and their results were also contested with arguments based on auditory processing of phasic information in the JO nerves of *Cx. quinquefasciatus* (Warren et al., 2009). However, the image channel resulting from the non-linear vibration of the antennae from the sound of the two sexes was shown to reinforce the hearing sensitivity of males close to/slightly above the frequency of the female’s second harmonic in electrophysiological measurements in *Cx. pipiens pipiens* (Lapshin, 2012) and *Ae. communis* (Lapshin and Vorontsov, 2021). The results of our behavioural assay suggest that this reinforcement is negligible in practice, at least in *An. coluzzii.*

The limitation of our stimulus recording approach is to be found in the long distances between the microphone and the single female (0.7±0.2 m), which induced a low signal-to-noise ratio of 1.7, despite noise filtering below the first harmonic and above the second harmonic (against a ratio of ~48 for the 2-harmonic 1-female stimulus; if considering the noise level as the noise floor between the 2 harmonics, using the Matlab function *snr*). Indeed, because of these different signal-to-noise ratios, the 2-harmonic 1-female stimulus can be seen as a frequency band of noise (ranging from the first to the second harmonic frequencies) instead of a true 2-harmonic sound.

However, this noise asymmetry between the two stimuli also shows that males are not fundamentally disturbed by noise; the noisiest stimulus (2-harmonic 1-female) induced as much response as the least noisy stimulus (2-harmonic 30-female). Wishart and Riordan (1959) found that female sound (500 Hz) is still an attractant to males, with at least up to 10 dB of noise above the signal sound level for *Ae. aegypti* males, but was not an attractant on the next tested step of 20 dB of noise above the signal level. The noise was composed of the superposition of sine waves of 100, 156, and 282 Hz plus square waves of 933, 1840, and 4130 Hz, which probably did not create as much noise around the female sound frequency as in our case. The hearing mechanism based on antennal distortion products uses the loud wingbeat frequency of the listener to amplify the nearby, but possibly quiet, wingbeat frequency of a potential mate (Lapshin 2012). By changing its own wingbeat-frequency, it is possible to change the distortion product frequency elicited by the nearby flying mate, which, theoretically, may help detect very faint harmonics against a relatively high level of background noise, especially when this noise is limited to the frequency band between the two harmonics.

### Constant sound *vs* ‘natural’ sound

Constant sound and female pre-recorded flight-tones have been known to trigger a response in mosquitoes for a long time (Roth, 1948; Kahn and Offenhauser, 1949). However, to our knowledge, no comparisons has been formally analysed between pre-recorded sounds and constant sound of the same frequency. Our results in Subset B show that the 1-harmonic constant sound behaves somewhat like a supernormal stimulus (for the wing-beat frequency response-parameter) compared to a 1-harmonic natural sound, at least at 26 dB SVL. Furthermore, Subset A allows us to conclude that males respond as much to 1-harmonic ‘natural’ sound as to 2-harmonic ‘natural’ sound. By combining results from Subset A and B, we deduce that mosquitoes hear natural sound as well as pure sound. This means that the information carried in the sound that elicits a male response is mostly the mean wingbeat frequency. A proper study could be carried out 1) with 1-harmonic constant sounds to control the sound level better than with pre-recorded sounds, and 2) by using larger ranges of frequencies and sound levels than in the present study, i.e., we would need to conduct a ‘behavioural audiogram’.

### Monitoring SVL from SPL measurements

Many studies report hearing thresholds based on SPL, which is a physical quantity that mosquitoes do not detect. We also monitored sound level with SPL, but we fulfilled the experimental conditions to provide equivalence between SPL and SVL, which mosquitoes do detect (see Methods section). Some studies have referred to SPL values as hearing thresholds, even though the equivalence conditions were not fulfilled or were unknown. Wishart and Riordan (1959) estimated that *Ae. aegypti* responds to a sound of approximately 20 dB SPL from experiments involving 30cm-side netting cages and sound stimuli presented through a diffuse speaker held against the cage netting. However, mosquitoes could be located a few centimetres from the loudspeaker, where SPL and SVL are not equivalent at this distance, i.e., when SPL would not be a good physical quantity to describe what the mosquito auditory organs are exposed to. Another example is Belton *et al.* (1961); a response threshold in the Johnston’s organs of male *Ae. aegypti* was measured to be between 0 and 10 dB SPL; SPL to SVL using a formulae that assumed far-field condition (without stating so, though). Unfortunately, the study did not provide enough details of the experimental setup to know the distance between the pressure microphone and the loudspeaker; thus, the thresholds were probably inaccurate. More recently, Dou *et al.* (2021) put their loudspeaker at 2.5 cm against their 30cm-side cage to measure the response of mosquitoes and monitored the sound level with an SPL meter in the middle of the cage. They measured flight response to sound in *Ae. aegypti* females for the first time, from a threshold of 79 dB SPL, which could be far more in terms of SPL since it was measured in the middle of the cage and mosquitoes were free to move along the cage’s sides, near the loudspeaker. In addition, SVL may have been far greater than SPL at this distance from the speaker. Taken together, SVLs probably do not occur with ecologically-relevant sounds, however, this could be used to inform the design of sound traps or reveal unknown auditory mechanisms.

## Acknowledgments

Natalie Morley (insect rearing), Stephen Young (discussion about statistics; design of the synchronization electronic tool; design of high-pass frequency electronics to adjust the dimming LED system).

## Data Availability

Raw response files (sound and tracked flight trajectories) are available at: https://doi.org/10.5061/dryad.9cnp5hqhj (Feugère et al., 2021a). Sound stimuli (Audio S1 to S4), as well as the original sound recordings (Audio S5, Audio S6) are available as supplementary material. Custom audio–video code for parameter-extraction and audio– video synchronization (modified Matlab files from https://doi.org/10.17632/hn3nv7wxpk.3 (Feugère, 2020)), custom statistics code for data analysis and figure plot (R files) and dataset (Text files) are available at https://doi.org/10.17632/6w5jttwkj8.2 (Feugère, 2021).

## Author contributions

Conceptualization LF, GG; Methodology LF, GG; Software LF; Formal Analysis LF; Investigation LF; Data Curation LF; Writing – Original Draft LF; Writing – Review & Editing OR, GG; Visualization LF; Supervision GG, OR; Funding Acquisition OR, GG, LF.

