## Supplementary Material for "Behavioural analysis of swarming mosquitoes reveals higher hearing sensitivity than previously measured with electrophysiology methods"

**TABLE OF CONTENT**

**Supplementary figures**

**Fig. S1.** Calibrated sound-level measurements of played-back sounds and background noise.
**Fig. S2.** Examples of *An. coluzzii* male response to the loudest four types of sound stimuli.
**Fig. S3.** Quantile-quantile plots on reduced model residuals.

**Supplementary tables**

**Table S1.** Filter characteristics applied to female sound recordings.
**Table S2.** Statistical tests on model null-hypothesis.

**Other supplementary files**

**Audio S1**
**Audio S2**
**Audio S3**
**Audio S4**
**Audio S5**
**Audio S6**
**Movie S1**

### Supplementary figures

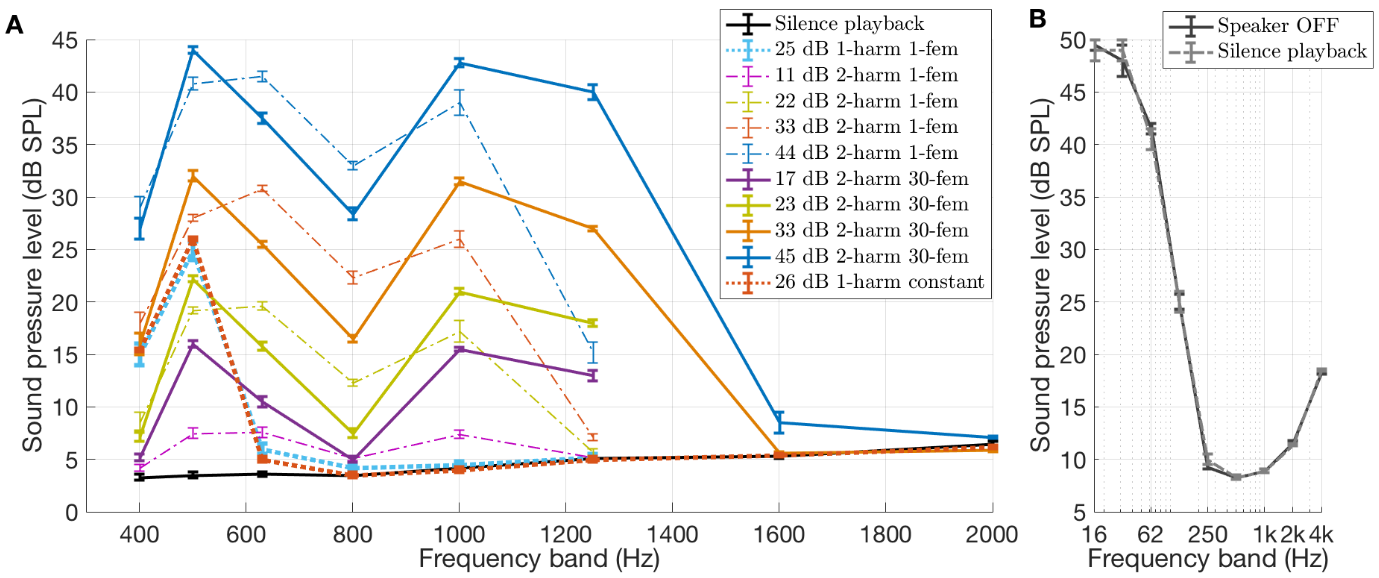


**Fig. S1. Calibrated sound-level measurements of played-back sounds and background noise.** SPLs (ref 20 µPa) were measured at the *Anopheles coluzzii* swarming position 1) to estimate the sound level received by the tested mosquito and 2) to know the background noise level of the sound-proof chamber. The X-axis values represent the central frequency $f_c$ of the octave or 1/3-octave band filters. Each filter has a lower limit of $2^{-1/6}f_c$ and an upper limit of $2^{1/6}f_c$. For example, X=800 Hz represents the sound pressure level between 713 Hz and 898 Hz. The Y-axis error-bar represents maximum and minimum values measured during stimulus duration under a time constant of 1 s (slow mode, according to IEC 61672-1: 2002). SPLs are equal to SVLs in our setup (see Method section 'Monitoring SVL from SPL measurements').

**(A)** SPL measurements of female sound stimuli as a function of frequency, limited to a frequency range audible to *An. coluzzii* males (Warren et al., 2009), and plotted at 1/3 octave steps reveals first and second harmonic of the female wingbeat sound. Black dashed line shows sound level when playing-back 'silent' (i.e., just speaker noise), with the same settings as during the experiment. The four coloured solid lines correspond to the sound levels at the mean male's 'XYZ' position during play-back of the female stimuli (related to four sound levels). Dashed lines represent the same, but for the 1-female sound-stimuli.

**(B)** SPL measurements in the sound-proof chamber without playback and with 'silence' playback, along octave bands from 16 Hz to 4 kHz. The black dashed line shows the sound level when playing-back silence (i.e., speaker noise), while the plain black line corresponds to when the speaker is off (i.e. showing noise floor of sound-proof room). Both are the same, i.e., they show that the speaker has a very low noise level enabling us to playback low level sounds.

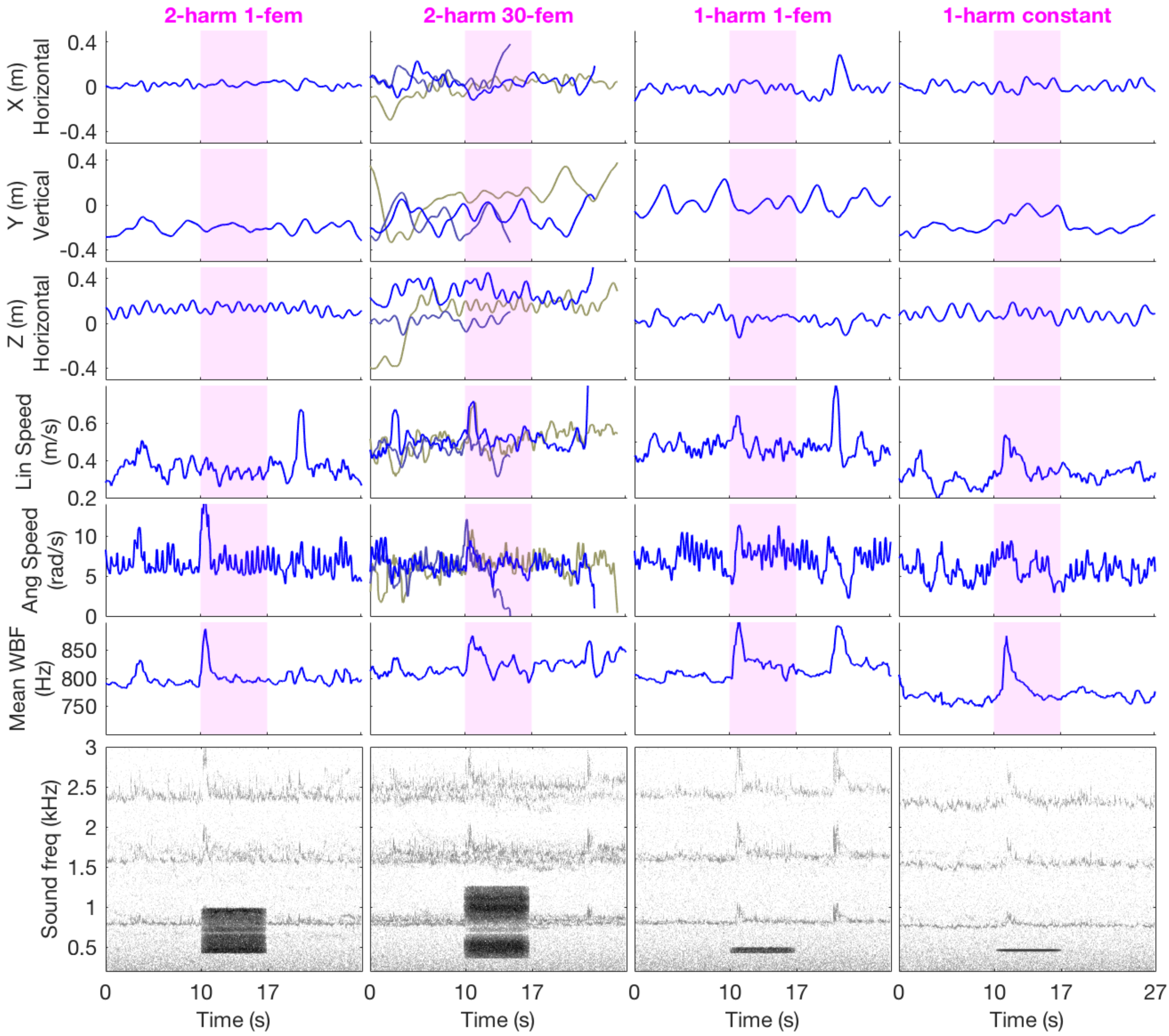


**Fig. S2. Examples of *An. coluzzii* male response to the loudest four types of sound stimuli.** Male flight-characteristics and wingbeat-frequencies (blue) before, during and after playback of female sound stimuli (red rectangle). Each column corresponds to a different type of sound stimulus (from left to right): 2-harmonic 1-female (Audio S1), 2-harmonic 30-female (Audio S2), 1-harmonic 1-female (Audio S3), 1-harmonic constant (Audio S4) . First five rows show flight parameters (relative X,Y and Z positions, plus linear and angular flight speeds). 'Z' dimension represents relative distance to the speaker (located 0.9 m from Z=0). Row before last shows mean wingbeat frequency over all present males (WBF), while the flight positions and dynamics corresponds to those of each present mosquito, hence the multiple lines on second column where 3 mosquitoes were present in this recording. Last row shows the spectrogram of sound recordings before, during and after the sound stimulus; the colour gradient represents the sound level given a frequency and a time (the darker the colour, the louder the frequency). See Fig. 2 for the spectrum of each sound-stimulus type.

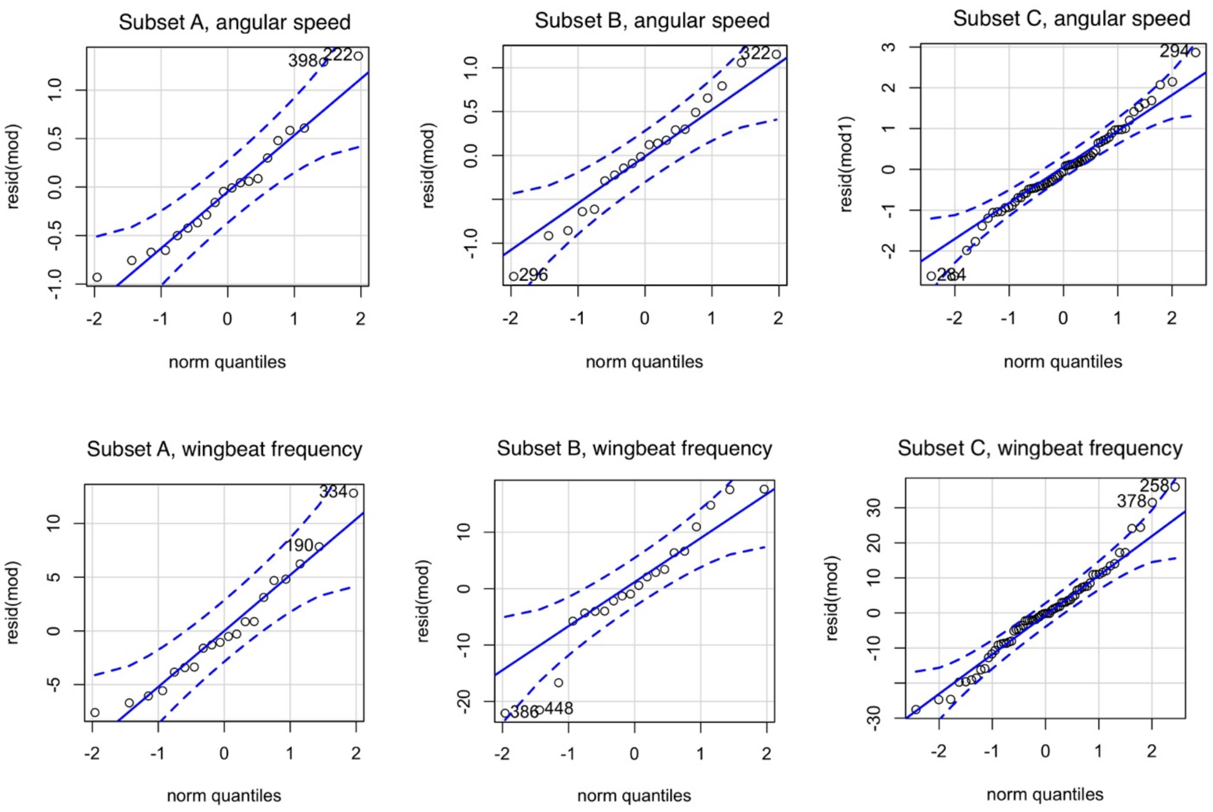


**Fig. S3. Quantile-quantile plots on reduced model residuals**. Qqplots were computed with the R *qqplot* function. See Table S2 for associated statistical tests.

### Supplementary tables

| Stimulus | FIR Filter type | Cut-off frequency (Hz) | Bandpass frequency (Hz) | Stop-band attenuation (dB) | Frequency (Hz) |
|---|---|---|---|---|---|
| 2-harmonic 30-female (Audio S3) | Highpass | 356 | 366 | 100 | / |
| | Lowpass | 1271 | 1261 | 100 | / |
| 2-harmonic 1-female (Audio S4) | Highpass | 422 | 432 | 100 | / |
| | Lowpass | 994 | 984 | 100 | / |
| 1-harmonic 1-female (Audio S5) | Highpass | 422 | 432 | 100 | / |
| | Lowpass | 497 | 487 | 100 | / |
| 1-harmonic constant (Audio S6) | / | / | / | / | 459 |

**Table S1. Filter characteristics applied to female sound recordings.** Filters were applied on sound recordings using the Matlab function *designfilt* with the parameters shown in the table, at a sampling rate of 8 kHz. The 1-harmonic constant sound was a single frequency/amplitude sound. For sound levels, see Table 1.

| | | Fligner-Killeen test (absence of variance homogeneity) | | | Shapiro-Wilk tests (absence of model residual normality) | |
|---|---|---|---|---|---|---|
| **Subset** | **Variable** | $\chi^2$ | **df** | **p-value** | $\chi^2$ | **p-value** |
| **A** | **AngSpeed** | 2.5 | 1 | 0.11 | 0.92 | 0.098 |
| | **WBF** | 0.0098 | 1 | 0.92 | 0.95 | 0.45 |
| **B** | **AngSpeed** | 0.0067 | 1 | 0.95 | 0.98 | 0.98 |
| | **WBF** | 1.2 | 1 | 0.27 | 0.94 | 0.26 |
| **C** | **AngSpeed** | 9.4 | 6 | 0.15 | 0.98 | 0.48 |
| | **LambertW-transformed WBF** | 8.8 | 6 | 0.19 | 0.98 | 0.30 |

**Table S2. Statistical tests on model null-hypothesis.** Tests were performed on reduced models for each subset and each extracted parameter **(**WBF: upper-quartile wingbeat frequency difference; AngSpeed: upper-quartile angular-speed difference) in R.

### Other supplementary files

**Audio S1 (AudioS3.mp3).** 2-harmonic 1-female sound stimulus (7 s).

**Audio S2 (AudioS4.mp3).** 2-harmonic 30-female sound stimulus (7 s).

**Audio S3 (AudioS5.mp3).** 1-harmonic 1-female sound stimulus (7 s).

**Audio S4 (AudioS6.mp3).** 1-harmonic constant sound stimulus (7 s).

**Audio S5 (AudioS1.mp3)**. Original sound recording of the 1-female *An. coluzzii* (7 s) before any filtering and level adjustment. Related to Fig. 2.

**Audio S6 (AudioS2.mp3)**. Original sound recording of the 30-female *An. coluzzii* (7 s) before any filtering and level adjustment. Related to Fig. 2.

**Movie S1 (MovieS1.mp4)**. Audio-video recording of a *An. coluzzii* male exposed to the loudest *An. coluzzii* 2-harmonic 1-female sound (10-s silence + 7-s sound exposition + 10-s silence). Related to Fig. 4 A.
